# The Low-Cost, Semi-Automated Shifter Microscope Stage Transforms Speed and Robustness of Manual Protein Crystal Harvesting

**DOI:** 10.1101/2019.12.20.875674

**Authors:** Nathan David Wright, Patrick Collins, Romain Talon, Elliot Nelson, Lizbé Koekemoer, Mingda Ye, Radosław Nowak, Joseph Newman, Jia Tsing Ng, Nick Mitrovich, Helton Wiggers, Frank von Delft

## Abstract

Despite the tremendous success of x-ray cryocrystallography over recent decades, the transfer of crystals from the drops where they grow to diffractometer sample mounts, remains a manual process in almost all laboratories. Here we describe the Shifter, a semi-automated microscope stage that offers an accessible and scalable approach to crystal mounting that exploits on the strengths of both humans and machines. The Shifter control software manoeuvres sample drops beneath a hole in a clear protective cover, for human mounting under a microscope. By allowing complete removal of film seals the tedium of cutting or removing the seal is eliminated. The control software also automatically captures experimental annotations for uploading to the user’s data repository, removing the overhead of manual documentation. The Shifter facilitates mounting rates of 100-240 crystals per hour, in a more controlled process than manual mounting, which greatly extends the lifetime of drops and thus allows for a dramatic increase in the number of crystals retrievable from any given drop, without loss of X-ray diffraction quality. In 2015 the first in a series of three Shifter devices was deployed as part of the XChem fragment screening facility at Diamond Light Source (DLS), where they have since facilitated the mounting of over 100,000 crystals. The Shifter was engineered to be simple, allowing for a low-cost device to be commercialised and thus potentially transformative as many research initiatives as possible.

**Synopsis:** A motorised X/Y microscope stage is presented that combines human fine motor control with machine automation and automated experiment documentation, to transform productivity in protein crystal harvesting.

## 1. Introduction

Since the 2000s, Macromolecular Crystallography (MX) has undergone a revolution in productivity, to become a high-throughput technique, thanks in large part to machine automation. Nanolitre-scale liquid handlers and robotic microplate imagers (Kuhn, Wilson, Patch, & Stevens, 2002; Stevens, 2000) are common in many laboratories. As is access to bright X-ray sources, high-speed X-ray detectors, and cryogenic sample changers that allow complete X-ray datasets to be measured in less than a minute (Bowler et al., 2015; Grimes et al., 2018). The notable exception to this trend in automation has been in the transfer of protein crystals from the crystallisation drop, usually in a microplate, to the sample mounts where they are stored for later X-ray diffractometry.

In most laboratories this harvesting step remains the same delicate, labour-intensive, manual process that it was at the advent of cryocrystallograpy (Garman & Schneider, 1997). One strategy to eliminate the mounting bottleneck has been to avoid the need for transfer entirely, by developing *in situ* diffraction techniques (Bingel-Erlenmeyer et al., 2011; Michalska et al., 2015; Soliman, Warkentin, Apker, & Thorne, 2011). Other approaches to *ex situ* screening have tried to design human out of the process, via novel harvesting techniques, or by reproducing the human mounting technique with advanced robotics (Cipriani et al., 2012; Deller & Rupp, 2014; Viola et al., 2011; Viola, Carman, Walsh, Frankel, & Rupp, 2007). Nevertheless, no affordable and scalable solution to the overall bottleneck of crystal transfer has emerged, and the systematic inefficiency of the harvesting step remains.

Crystal harvesting is a task that exists as part of a broader experimental workflow. As a skill manual mounting is easy to learn, but difficult to master; minimising experimental variability between crystals and maximising mounting productivity requires simultaneous management of multiple challenges: fine movements and sensory input to manipulative crystals gently; awareness of changing drop conditions; organisation of multiple sample plates; and thorough data management. Unsurprisingly, manual mounting of crystals presents a source of experimental variability and sample loss, as well as a process bottleneck in the MX workflow.

As experimental throughput is increased, manual mounting and experimental documentation become limiting. For instance, in the 11 years from April 2004 to 2015, the 1718 structures released to the Protein Data Bank by the Structural Genomics Consortium (SGC) involved mounting 48,373 crystals by hand, an average of 4,397 crystals per year. The time taken by manual mounting and data management at this scale places an artificial limit on the kinds of experiments that can be done. Crystal fragment screening, which requires 100s to 1000s of crystals per target, is another use-case that is limited by the absence of a solution to crystal transfer.

No solution to the mounting bottle-neck has yet achieved wide-spread adoption, and this may be for a variety of reasons: incompatibility with existing practices, high initial cost, engineering support burden, or lack of commercial availability. Perhaps the most fundamental obstacle to fully automating x-ray crystallography are the technical difficulties crystal harvesting presents. Locating crystals requires high resolution imaging in all three spatial axes (x,y,z), as well as in the temporal axis. Crystals can move by as much as 15µm/s due to fluid dynamical effects (Savino & Monti, 1996), and will be further disturbed during mounting (Read, P. & Meyer, 2000). Whilst most crystallography labs will have a stereoscopic microscope, compatible with the Shifter, for manual mounting stereoscopic digital imaging systems do not seem to offer sufficient z-axis resolution at the temporal resolution required (Kwon et al., 2010; Pei, Xu, Zhu, & Wang, 2012; Dean et al., 2017; Štolc, Soukup, Holländer, & Huber-Mörk, 2014; Levoy, Ng, Adams, Footer, & Horowitz, 2006). Furthermore machine identification of protein crystals from digital images has also proven exceptionally difficult (Liu, Freund, & Spraggon, 2008). There are significant variations between imaging conditions and crystal morphologies are highly varied; they are often small (c.10-75µm), colourless, display poor optical contrast with the surrounding droplet (Nollert, 2003), and can be obscured by other droplet features. A significant body of research has accumulated since the emergence of high-throughput MX, on the problem of accurately identifying crystals (Ng, Dekker, Kroemer, Osborne, & von Delft, 2014), yet solutions have been only partial.

To manipulate fragile protein crystals requires fine movements and rapid sensory feedback. These are all complex and costly engineering problems to solve. Fully-automated crystal mounting, compatible with existing experimental practice, and accessible to the wider community may therefore, be some years off (Deller & Rupp, 2014).

Instead we designed the Shifter which exploits human aptitude combined with minimal machine automation and software workflow tools. We show here that whilst such a simple solution to these engineering challenges is nonetheless sufficient to address the productivity gap of harvesting. The Shifter allows the human mounter to focus their adequate visual acuity, dexterity, and sensory feedback to specialise in mounting without distractions. By automating only those aspects of mounting humans are less good at, such as the repetitive organisational tasks that make mounting slow and unreliable, a division of labour is achieved that greatly improves productivity, whilst avoiding engineering complexity.

## 2. Methods

### 2.1. Basic principle of operation

The Shifter is an X-Y stage allowing one or two microplates to be loaded into the Shifter via a load port in the enclosure lid. Immediately before loading microplates into the Shifter the film seals are completely removed. The mounter uses a touch screen PC to request mounting targets to a hole in the lid, at the microscope optical axis, where the crystals are harvested by the human, in the normal

**Figure 1.**
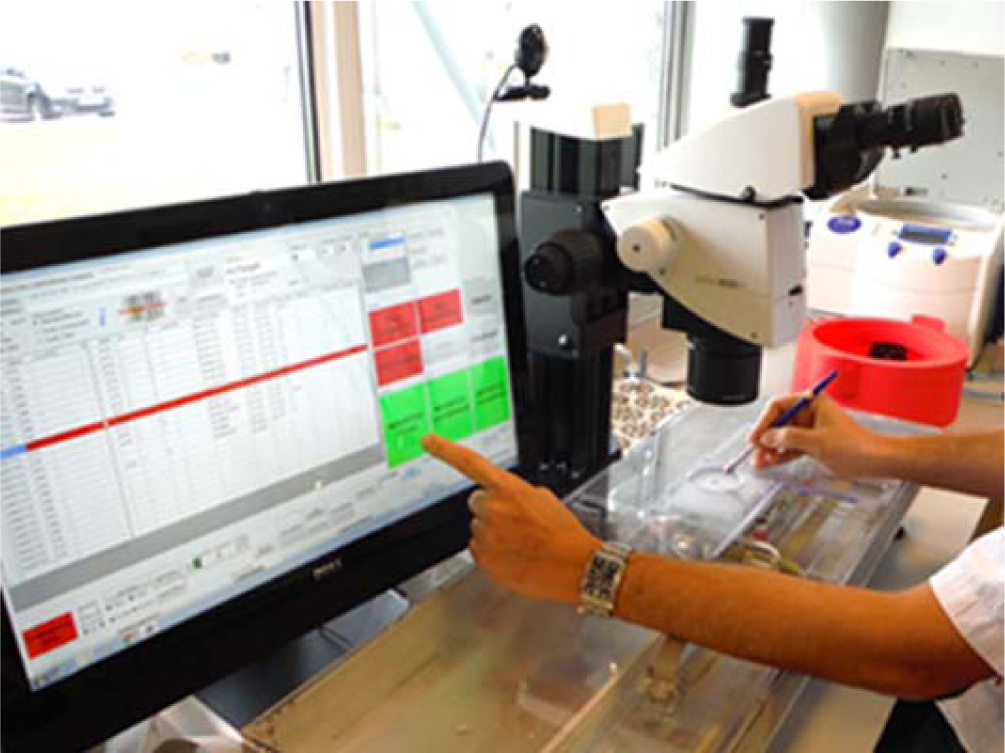
The Shifter deployment at XChem: The operator is mounting through the mounting aperture, whilst operating the GUI with the free hand. The interface manoeuvers samples within the device enclosure, whilst automatically completing experimental annotations and documentation.

### 2.2. Description of the device hardware

The Shifter enclosure (*Figure 2*(a,b)) is metal (1), with a clear plastic lid (2) that has a large port for loading plates (3), and three small ports for access to loaded plates: the Mounting Aperture, concentric with the optical axis of the microscope (4.1), and expansion ports (4.2). The Shifter is installed at a mounting microscope (*Figure 2*(a)). Microplates, loaded with the seals completely remove, are manoeuvred in the X and Y directions by means of stepper motors and toothed belts, with positional feedback from linear encoders (Spectra Symbol, Salt Lake City, UT). Any part of the left or right microplate can be positioned at the Mounting Aperture, for human mounting, or at the expansion ports, whilst every other part of the microplates remain sealed.

The x-axis carriage (5) (*Figure 2*(c)) moves in tracks on the enclosure base on low-friction polymer linear slide bearings (igus GmbH, Cologne, Germany). The y-axis carriage (6) travels in guide tracks on top of the x-axis carriage using a similar method of transmission. Two independent plate carriers (7) hold one microplate (8) each, and move freely in the z-axis. This low-cost and fit-for-purpose stage construction contrasts with typical motorised microscope stage construction, which often use high-cost, precision made components that also require tighter tolerances in the manufacture of the assemblies to which they are mounted.

**Figure 2.**
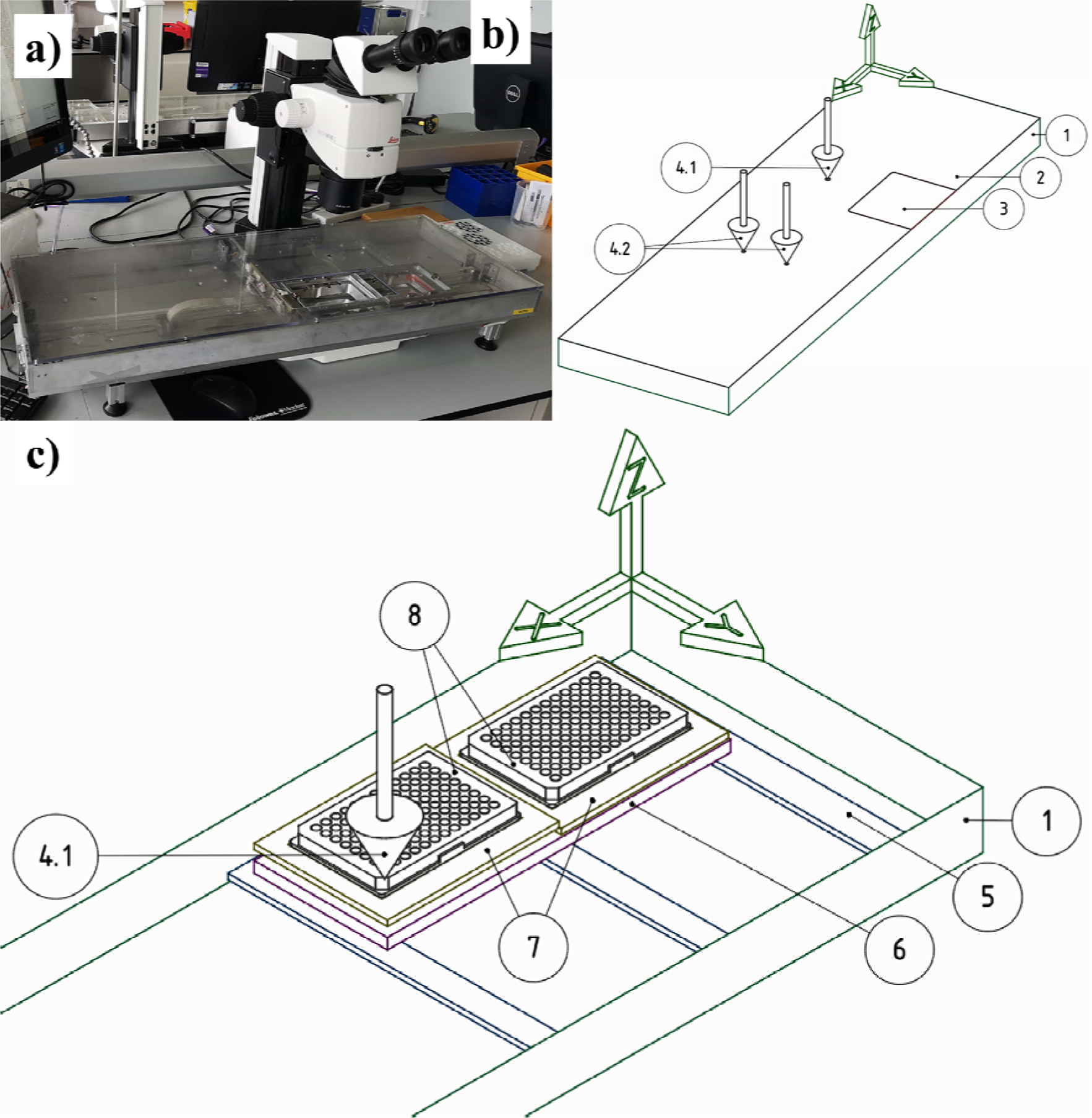
**(a)** The Shifter as installed at a mounting microscope. **(b)** A simplified drawing of the main elements of the Shifter enclosure (c.90cm x 30cm x 6cm). **(c)** A simplified drawing of the microplates in relation to the x-axis carriage, the y-axis carriage, and the z-axis mechanism

#### 2.2.1. The mounting aperture and expansion ports

The plate access ports have clearance envelopes around them such that any part of the microplate can be placed under that port whilst all other parts of the microplates remain sealed (**Figure *3***). The mounting aperture was profiled to provide protection to wells adjacent to the well in-use, while allowing a full range of mounting angles (**Figure *3*a**). The two expansion ports are to accommodate additional functionality.

**Figure 3.**
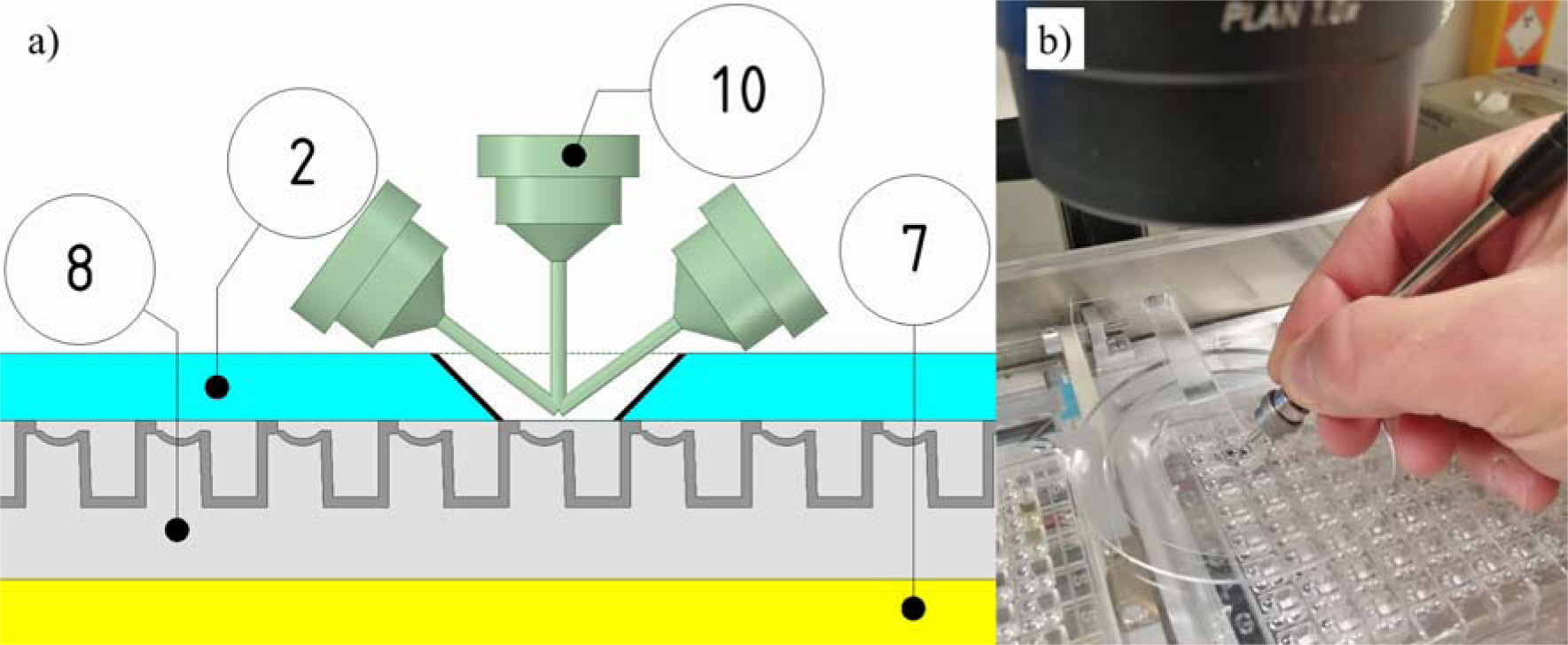
Mounting Aperture; access to the microplate (8), on its carrier (7), through a hole in the enclosure lid (2). **a)** Cross-section showing a range of mounting angles for the pin (10), through mounting aperture. **b)** A pin being used for mounting through the Mounting Aperture.

#### 2.2.2. Z-axis movement: protecting plates without film seals

When the stage is in motion, the microplates are pulled away from the enclosure lid, against the force of supporting springs, by two voice coil electromagnets (MotiCont, Los Angeles, CA) on each microplate holder (Figure 4). The voice coil motors are de-energised at the end of the move, releasing the microplates, and reforming the contact between plate and lid. The stage’s two microplate holders can be adjusted separately and setup for different microplate heights and masses.

**Figure 4.**
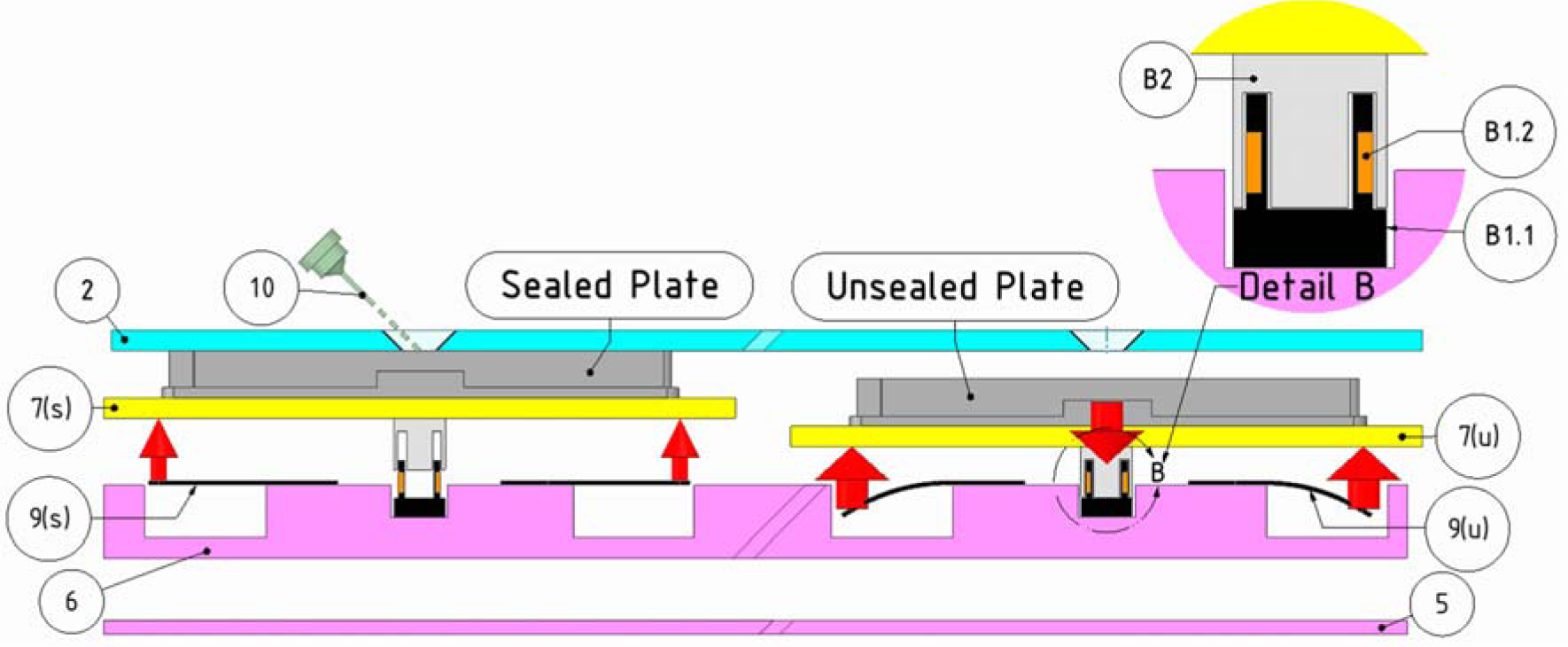
The Shifter mechanism for sealing microplates with the film seal completely removed: (10) Crystal mount pin, (2) acrylic lid, (6) y-axis carriage, (5) enclosure base. LEFT: The plate shown in its carrier (7(s)), in the sealed position between moves. It is supported by flat springs (9(s)). RIGHT: during a move the plate carrier (7(u)) is pulled down against the force of the supporting springs (9(u)) by electromagnets (Detail B). **Detail B**: Voice-coil electromagnets comprising coil holder (B1.1), coil (B1.2), and permanent magnet cap (B2). The red arrows indicate the magnitude and direction of forces applied to the microplate holders.

### 2.3. Stage feedback and control

#### 2.3.1. Sensor and Control Electronics Modules

Electronics sensing and control modules from Phidgets Inc. (Calgary, Canada) were used as they can be connected directly to a PC via USB without intermediate electronics. The vendor provides software drivers and libraries that support application development in a wide range of operating systems and programming languages. Software applications running on the PC integrate the individually addressable Phidgets modules programmatically. Having a variety of module options, which are easy to integrate, costs somewhat more compared to open-source alternatives, however we found that this convenience greatly accelerated prototyping.

We considered implementing continuous, smoothed motion for moving plates, known as ‘tool paths’, where a stage or tool follows every point of a predetermined route between locations of interest. We concluded the current Point-to-Point movements to be sufficient for crystal harvesting. Tool paths would require tightly coordinated multi-axis movements, synchronised at a low-level, electronically, which is not possible with Phidgets. Moving between points of interest on a microplate does not require defined tool paths, so we took the view that the additional development and expense this would have required, was not warranted.

#### 2.3.2. Stage moves and feedback coordination

Stepper motors are a common choice for positioning applications as they are inexpensive and are easily controlled programmatically, by requesting a given number of ‘steps’ of rotation. Open-Loop systems, where there is no positional feedback, are the simplest configuration to implement. However the looseness and flexibility in any drive chain leave it susceptible to losing position. Due to the mass of the moving parts, fast rates of acceleration, travel, and deceleration lead to missed motor steps and target overshoot.

In Open-Loop systems the problem of losing position can be resolved by using over-sized motors, able to cope with the forces involved. In the Shifter, space constraints limit motor size, therefore position encoders are needed to provide positional feedback. The stepper motors are therefore controlled in a Closed-loop configuration. Rotary encoders mounted on the motor driveshaft are commonly used for motor position feedback and missed step detection. However, the Shifter’s relatively low-cost construction means that there is significant slack in the transmission, between the motor shaft and the stage that rotary encoders would miss. Ultimately, only by tracking the position of a point fixed in relation to the microplate, can the position of a given drop be reliably determined. Thus we located the contact point for each axis’s encoder in the relevant stage carriage.

By using linear encoders the positional resolution of the stage is related not to the step size of the motors (25µM typical), but to the resolution (and linearity) of the encoders used. Whilst the resultant positional accuracy (0.1-0.15mm) is coarse for a typical X/Y stage (3.4), it is compensated for by human aptitude, thus engineering complexity is kept down, reducing costs.

### 2.4. Support and documented of diverse workflows

A prototype Graphical User Interface (GUI) was necessary to test the system. The GUI was developed as a Windows Form Application (.NET Framework) using the Microsoft Visual Studio integrated development environment (IDE), and was coded in C#.

A manual interface within the GUI provides casual microplate access; the user selects the microplate and subwell, on the touch screen computer, and the stage drives the requested location to the Mounting Aperture (Figure 5a). No experimental annotations are captured in this mode.

**Figure 5.**
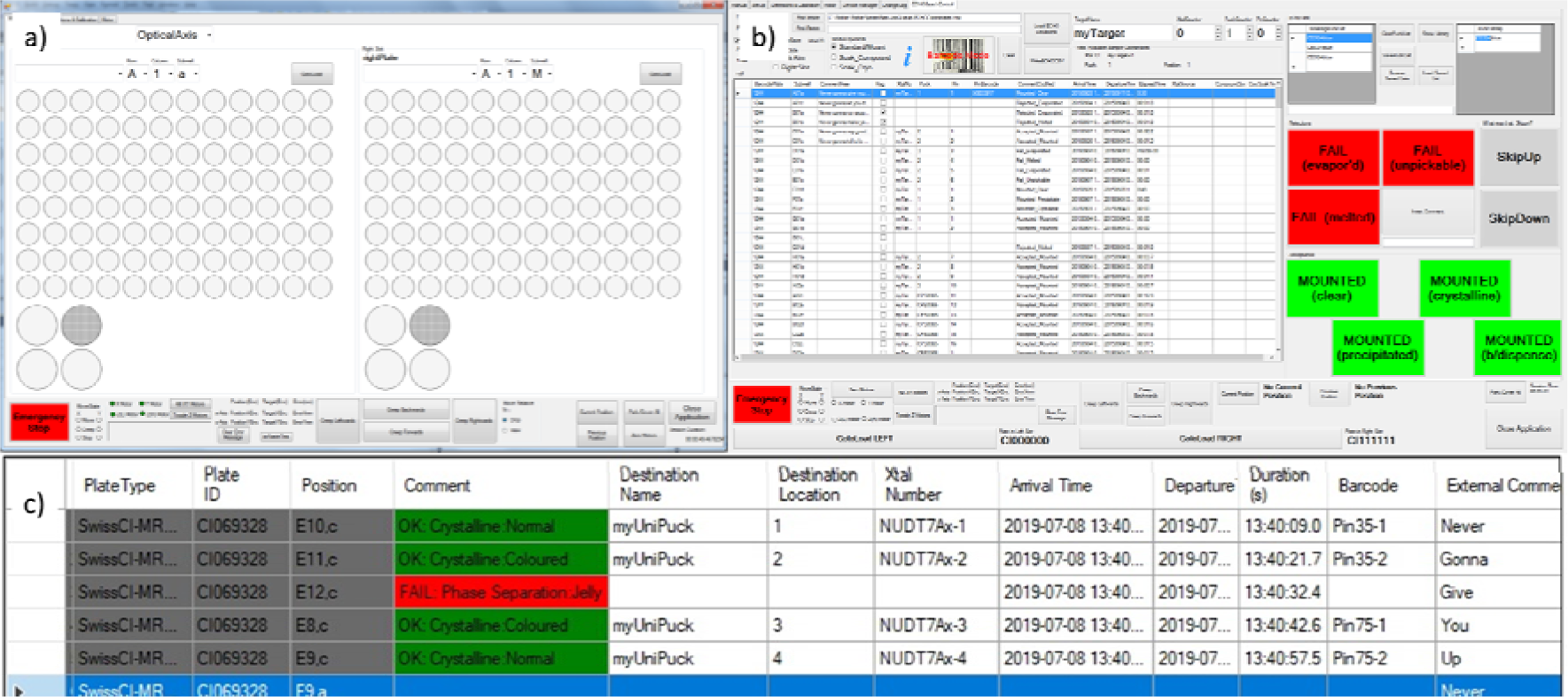
**a)** Undocumented Mode: a manual driving interface allows any subwell to be selected, and driven to the desired stage position. These moves are not documented. **b)** Guided crystal mounting interface; the table contains a picklist of the locations to be mounted from, and touch-screen buttons for driving the stage to the next location (shown in red/green). **c)** A close-up of the picklist table with columns for tracking data and experimental outcomes which are populated automatically in real-time

Where locations of interest are known in advance they are imported into the Shifter GUI from a .CSV (comma separated variable) file. The imported file can contain locations from any number of plates or experiments. The mounter proceeds down their list of targets using large touch-screen buttons, and that location is moved into position. These configurable buttons allow the user to conveniently control the workflow with the non-mounting hand. In documenting mode these buttons also automatically capture the experimental outcome, alongside the sample tracking data generated. This data is saved for export as a .CSV file that can be imported back into the user’s workflow *e.g.* a database or Laboratory Information Management System (LIMS). Correlations between experiment outcomes and protocols can then be routinely identified, a capability not previously available. This is the principle data source for the statistical analyses in this study. To demonstrate generality two experimental protocols were implemented, which provide automatic experimental documentation:

#### 2.4.1. Example 1: Simple guided mounting

In simple mounting mode, one or two microplates are loaded at a time, from the list locations imported into the interface (Figure 5b). As users navigate down the work list, the relevant plate and location is physically moved to the Mounting Aperture. When the loaded plates have been processed, the user is prompted to load the next plates in the series. Annotations are saved as previously described.

#### 2.4.2. Example 2: Fragment soak

An ‘advanced mode’ facilitates cryo-protectant flash-soaking, and harvest-soak-retrieve compound soaking processes. Here the user operates a table of crystal source locations, and a second table of soak locations. Navigating between the two, source plate and destination plate locations are presented at the Mounting Aperture, where crystals are first mounted, transferred to the soak condition, and later retrieved. Fields in the user work list are automatically populated with tracking data linking unique crystal identities with soak conditions for export as a .csv file.

#### 2.4.3. Example 3: Adaptations to additional experimental protocols

Large-droplet-format hanging drop and sitting drop crystallisation experiments were enabled through specially designed and 3D-printed adaptors. These adaptors accommodate crystal systems presented on 18mm and 22mm round or square cover slides, or microbridges, such as those used with VDX™ (Hampton Research, Aliso Viejo, CA) or Linbro® (MP Biomedial, Santa Ana, CA) plates (S4, Figure 13 & Figure 14).

### 2.5. Validation experiments

The device was validated by assessing both user acceptance, and whether the engineered solution demonstrated usability, efficacy of the sealing solution, and mounting productivity gains. Firstly, the Shifter was deployed at the XChem facility, which was followed by experiments designed to demonstrate that an engineered solution to the mounting problem was superior to the existing manual practices (2.1-2.4.3).

For all of these experiments the Shifter was used to mount in documenting mode, where timestamps and experimental annotations are automatically generated, and exported as a .CSV file (2.4.1). These files were retrieved and used to calculate relative mounting rates, and total productivity as measured by total number of mounted crystals and x-ray diffraction datasets collected.

#### 2.5.1. Sample preservation without film seals

Microplates are usually with an adhesive film during storage, which must then be excised during mounting. The Shifter avoids manual film cutting using the sealing mechanism described in section (2.2.2, Figure 4). To evaluate evaporation after film removal a simple test was devised wherein 50nL droplets of 1.5M NaCl were deposited into a microplate and monitored for nucleation, as an analogue of droplet evaporation. Time to nucleation was measured for a droplet under the Shifter lid at the Mounting Aperture, with and without a proposed draft excluder (2.4.3), and a positive control of exposed droplets set on top of the enclosure lid (results in section 3.1).

#### 2.5.2. Relative Productivity: Assessment of mounting rate

A productivity base-line for mounting crystals was established by surveying SGC mounters about practices encountered in the community and expected mounting rates (S2, Table 2 and Table 3). Self-reported data from the six respondents was used to calculate mounting rates measured in crystals mounted per hour, or minutes required per mounted crystal.

Exported Shifter data .CSV files from XChem user sessions for the period September 2015 to January 2016 were aggregated, and analysed for patterns of behaviour from academic and industrial crystallographers.

Next a comparison was made between mounting rates using the manual mounting process (cut-and-reseal film) and the Shifter-assisted process, for a novice mounter (NDW) and an experienced crystallographer (PC), to test how mounters of different experience respond to the Shifter (study protein DACASA).

Finally a case-study was carried out to explore the burden associated with training and familiarisation of users to the Shifter. In this study a trainee was given brief instruction (c.10mins) on Shifter use, having had no previous crystal mounting experience, after which they mounted unsupervised, from a study protein (pnp2).

Results are discussed in section 3.2.

#### 2.5.3. Absolute Productivity: Quantity and quality of crystals retrieved using the Shifter

We measured absolute productivity by the number of crystals mounted, and by X-ray diffraction datasets collected. To test for any effect on absolute productivity from using the Shifter we set up microplates of NUDT7A with conditions know to give an abundance of crystals 35-75µm in size. Droplets were imaged and assessed for the presence of such crystals, easily accessible to a novice mounter (NDW) (Figure 8). Second choice target crystals (10-35µm in size, or those poorly accessible) were also documented as they might be a valuable data source in real situations. Microcrystals <10µm were not counted, as although they can be mounted using the Shifter, this is not typical for our current workflows. The microplates used have three subwells in each of the 96 well locations; when a well is unsealed, all three subwell drops in that well are exposed. For this experiment an initial subwell drop was chosen from a suitable well, and mounted from for as long as was possible. When all accessible crystals in the initial drop were mounted, the other drops in the exposed well were used. A cohort of five wells were mounted from using the traditional manual method of cutting and removing the seal (‘Manual’). A second cohort of wells was mounted with the microplate placed in the Shifter, but with the stage stationary between each mounting event (‘Shifter Stationary’). A third group were mounted from the Shifter with a stage move between mounts, to simulate a soaking step or similar (‘Shifter Moving’). The collected crystals were then evaluated to determine the x-ray diffraction limit. (Results are discussed in section 3.3).

#### 2.5.4. Further strategies for decreasing evaporation from samples

In addition to preventing drop drying decreasing dehydration in crystals during mounting has been reported to improve the reproducibility of unit-cell (Farley et al., 2014). In the absence of a universal used method for the humidification of samples during mounting, strategies at the SGC have included placing moisture sources around the mounting area, or directing the output of an ultrasonic humidifier onto the exposed droplet. These solutions can be exquisitely sensitive to disturbances in air currents within the mounting room, and in the case of ultrasonic humidifiers, generate an aerosol of water droplets that pools on the work area.

A system was developed control the mounting environment, between the microscope objective and the exposed drop using a draft excluding shield, and a low-cost humidifier built from commonly available parts (**Figure *6***)

**Figure 6.**
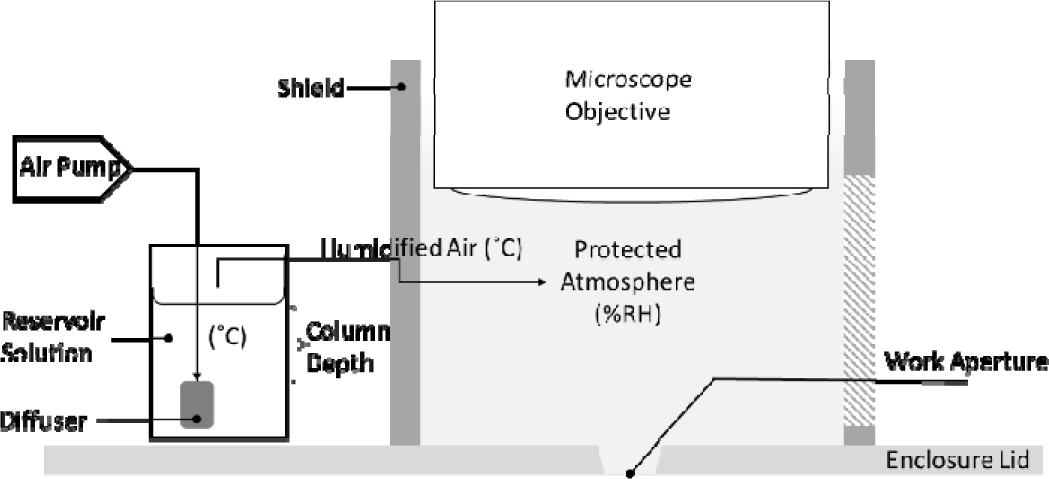
Schematic of shield and humidifier, made from an aquarium air pump, diffuser (air stone), silicone tubing, and a laboratory bottle fitted with a twin-port lid. The pump output (2×250 L/h) was connected to a single 19mm x 42mm rod-shaped diffuser.

For a given gas flow rate the simplest way to increase %RH is to increase the depth of the reservoir column above the diffuser (Al Ashry & Modrykamien, 2014). Evaporative losses make maintaining column depth problematic, so it should be fixed to ensure 100%RH across a range of conditions, then blended with dry air to the desired %RH. An approach similar to this was employed by Christopher Farley and colleagues in their work to improve the reproducibility of unit-cell parameters using a custom apparatus, to limit crystal dehydration during mounting. (Farley et al., 2014). (Results are discussed in 3.1, for data see S4).

#### 2.5.5. Effect of low-cost components on positional accuracy

We evaluated the effect of low-cost components (section 2.1) on system performance. Careful calibration was combined with modelled error fitting to compensate for a lack positional accuracy (S1, Figure 11a). Membrane potentiometer position sensors were used to encode the location of the stage in X and Y axes, because of their low cost and simplicity of installation. Resistance or voltage (as ratio of supply voltage), is measured across the encoder to determine the position of the stage axis. To relate encoder values to real-worlds coordinates scale tape is applied to the enclosure base along the x and y axes. A USB microscope is fixed to the stage so that crosshairs on the camera image overlay the scales (S1, Figure 11b). This allows encoder readings to be taken periodically along the millimetre scales, for the full length of the encoder. Live, real-world stage coordinates can thus be retrieved by converting encoder readings in real-time via a polynomial function fitted to the calibration data. Polynomial functions of increasing orders were trialled in order to find the optimal function for real-world coordinate elucidation (results are discussed in 3.4).

## 3. Results & Discussions

The development of the XChem facility for fragment screening in protein crystals in 2015 established as routine, a technique with a fundamentally higher demand for mounted crystals than the standard crystallography experiment; XChem has averaged 25,000 crystals per year in its first four years (Collins et al., 2017; Cox et al., 2016; Krojer et al., n.d., 2017; Mcewan et al., 2010; N. M. Pearce et al., 2017; von Delft, n.d.). The scale-up of XChem into an industry and academic facility has further accelerated demand for mounted crystals, and has only been possible through the successful development and deployment of the Shifter, and its associated increase in productivity. Beyond this we devised additional experiments to demonstrate vigorously how drop and crystal survival are improved.

### 3.1. Shifter plate sealing method dramatically reduces droplet evaporation

Microplates are loaded into the Shifter without any film seal (2.5.1), relying instead on the mating of the upper microplate surface with the under-side of the enclosure lid. We show that the Shifter greatly slows evaporation compared to film cutting-and-resealing, reducing stresses on the crystals and increasing the time mounters have to work on the drop before it dries.

Nucleation of aqueous NaCl occurs circa 6.1M under ambient conditions. From a starting concentration of 1.5M in 50nL drops this represents a loss of ¾ of the water, or approximately 38nL.

**Table 1.**
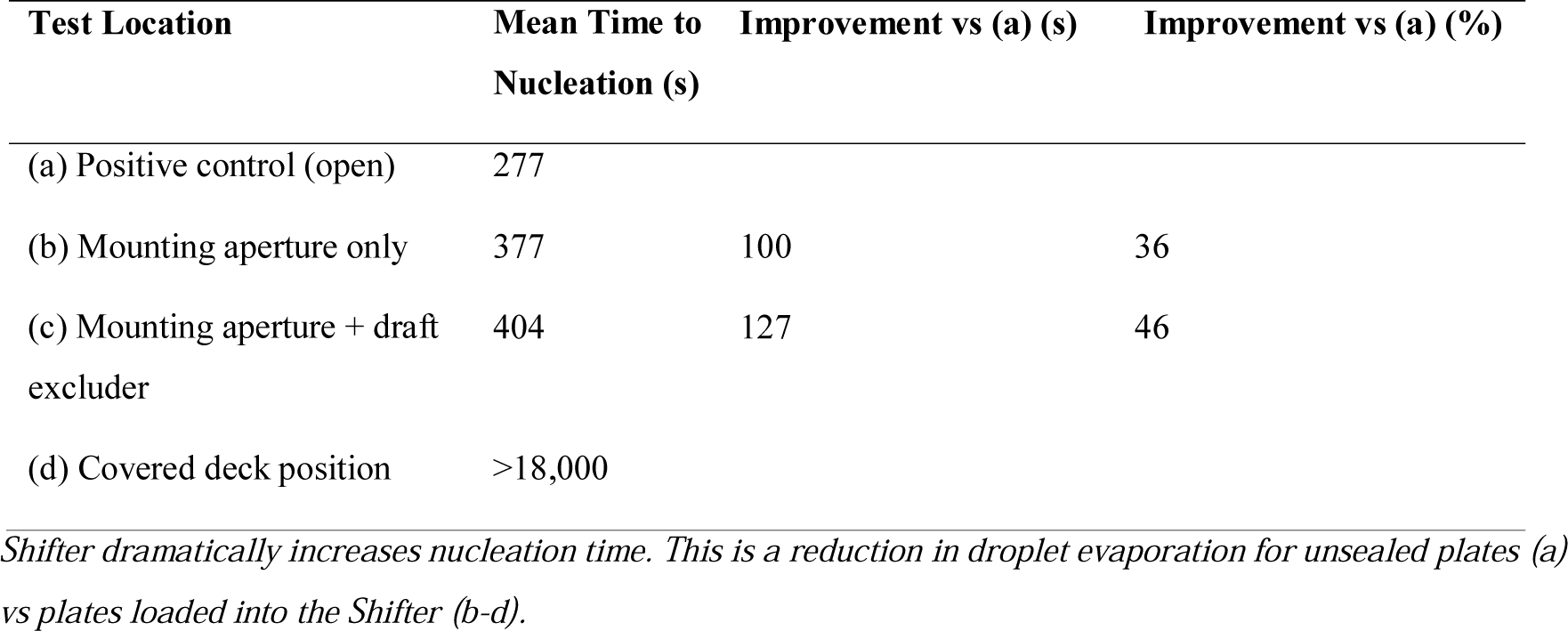

| Test Location | Mean Time to Nucleation (s) | Improvement vs (a) (s) | Improvement vs (a) (%) |
| --- | --- | --- | --- |
| (a) Positive control (open) | 277 |  |  |
| (b) Mounting aperture only | 377 | 100 | 36 |
| (c) Mounting aperture + draft excluder | 404 | 127 | 46 |
| (d) Covered deck position | >18,000 |  |  |

*Shifter dramatically increases nucleation time. This is a reduction in droplet evaporation for unsealed plates (a) vs plates loaded into the Shifter (b-d).*

### 3.2. Mounting Rate: Relative productivity is improved for all users

Self-reported data from a survey of six SGC crystallographers suggests a productivity baseline for manual mounting of 8 crystals/hour, when list generation, data entry, and time spent on all other aspects of manual mounting are included. Whilst mounting rates will be highly dependent on the mounter and the protein system in question, this productivity baseline is consistent with the authors’ experience. In the search for a solution to the mounting bottleneck it is significant that respondents estimate between a quarter and a half of mounting time is in reality spent on ancillary tasks (S2, Table 2) such as this represents a new avenue for process optimisation in mounting.

XChem users were able to mount 8271 crystals from at least 17 crystal systems using the Shifter, in the first four months of its deployment (Sep. 2015-Jan. 2016). Experimental outcomes automatically captured by the Shifter GUI, show 86% of mounts were judged to have been a ‘success’ by the user, with a mean mount time of 35 seconds (103 crystals/h), and a median of 30 seconds (120 crystals/h) (Figure 7b). This is in contrast to the productivity baseline estimate of 7 ½ minutes per mounted crystal.

To see if the Shifter could act as shortcut to the greater productivity that comes with being an experienced mounter, we compared mount durations between a novice (NDW) using the Shifter and an expert (PC) mounting manually, but found no difference in mount duration. This method was sensitive to a difference between novice and expert when both used the Shifter (p<0.000). We conclude that Shifter-assisted novices can become as productive as manual experts, but that the Shifter increases productivity for all levels of mounter.

In separate case-study of a ‘Shifter Trainee’, after 10 minutes of training, mean mounting rates of 75 crystals/h were seen initially, rising to 140 crystals/h after 5 hours of accumulated mounting time (Figure 7, a)). The starting productivity rate was an order of magnitude faster than the survey baseline and continue to get a lot quicker, at a rate of 22 crystals/h for each additional 100 crystals mounted (R^2^=0.75).

**Figure 7.**
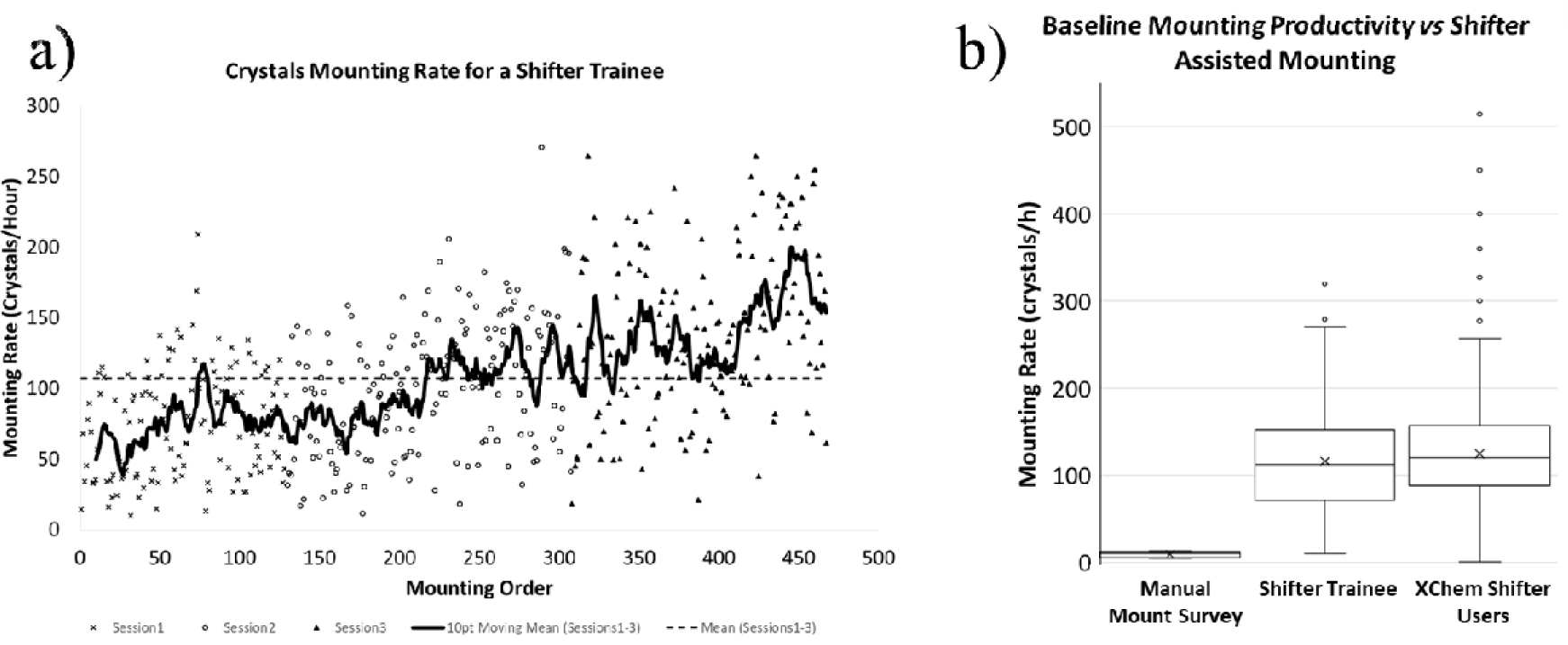
**a)** The Shifter Trainee quickly improved productivity. After c.250 crystals this novice mounter achieved a typical rate of 130 crystals/h; crystals/h=(seconds per hour)/mount duration(seconds). **b)** A comparison of mounting productivity: The results of the SGC estimated ‘Manual Mounting Survey’ rate (S2, Table 3), the ‘Shifter Trainee’, and data from ‘XChem Shifter Users’ (September 2015-January 2016) (S3, Figure 12). All data is calculated from automatically stored timestamps generated by the Shifter GUI. Mount duration is the difference in time between the requested drop arriving at the mounting aperture and the user requesting to leave it.

### 3.3. Absolute Productivity: Shifter allows mounting of more crystals that diffracted

In the experiment to quantify the effect of the Shifter on the absolute number of crystals mounted, and datasets collected (2.5.3) we saw that for the Manual wells, once the initial drop became unusable all adjacent drops in were also unusable. In the Shifter experiments wells the adjacent drops were still viable for mounting, and the diffraction limits for the crystals collected from those subwell drops weren’t significantly different to those crystals from initial drops (t-stat: 2.000, p=0.14).

**Figure 8.**
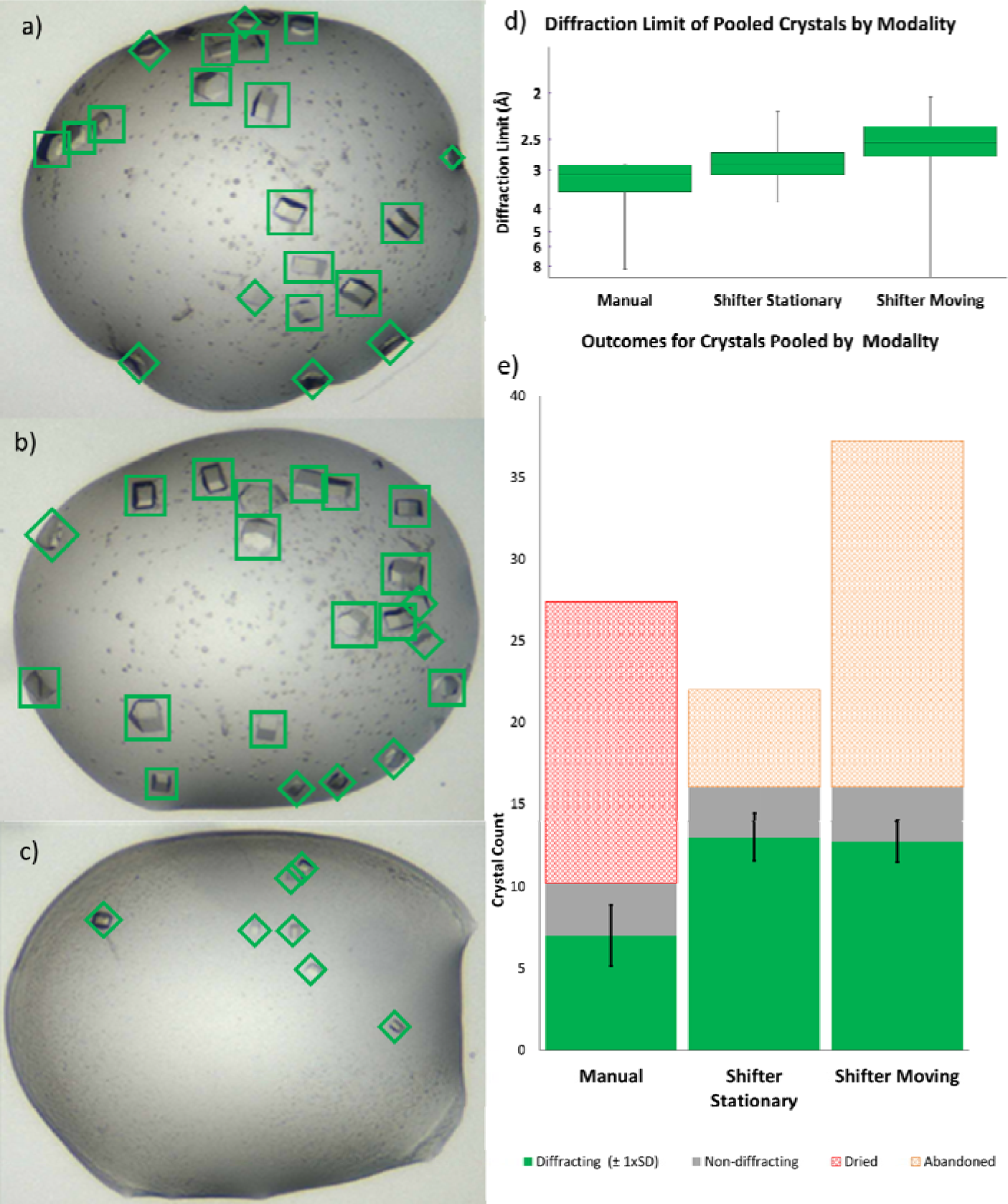
The Shifter enables more crystals (NUDT7A) to be mounted. **(a-c)** A well drops contains multiple crystals were identified from images based on the number of easily accessible 35-75µm crystals (green squares). 10-35 µm, or inaccessible crystals, were counted as second-choice targets, which might be of interest to under real conditions (green diamonds). Microcrystals were not counted. Images shown (a-c) relate to experiment 1 in the ‘Shifter Stationary’ series (Figure 10, middle pane). **d)** The X-ray limits of diffraction were similarly distributed for crystals mounted from Manual and Shifter experiments. **e)** Practical considerations limited us to one Uni-Puck per well (16 crystals). For Manual wells drops dried-out before reaching this limit (red hatching). For the Shifter wells this quota was met, necessitating that remaining viable crystals be abandoned (orange hatching).

Mounted crystals yielding a diffraction dataset for the Shifter experiments show a significant improvement in success-rate of mounting over the Manual process (Shifter Stationary, t-Crit. 1.9, p<0.001; Shifter Moving, t-Crit. 1.9, p<0.001). No significant difference exists between the two Shifter experiments (Figure 8d). When all drops from each cohort are pooled we also found an improvement in the high resolution limit for diffraction from the Manual crystals to the Shifter Stationary (t-Stat: 2.48, p0.018), and from Stationary to Shifter Moving (t-Stat: 2.47, p0.015). This suggests that stage movements provide a small additional improvement on crystal survival, on top of the highly significant improvement over the Manual process.

**Figure 9.**
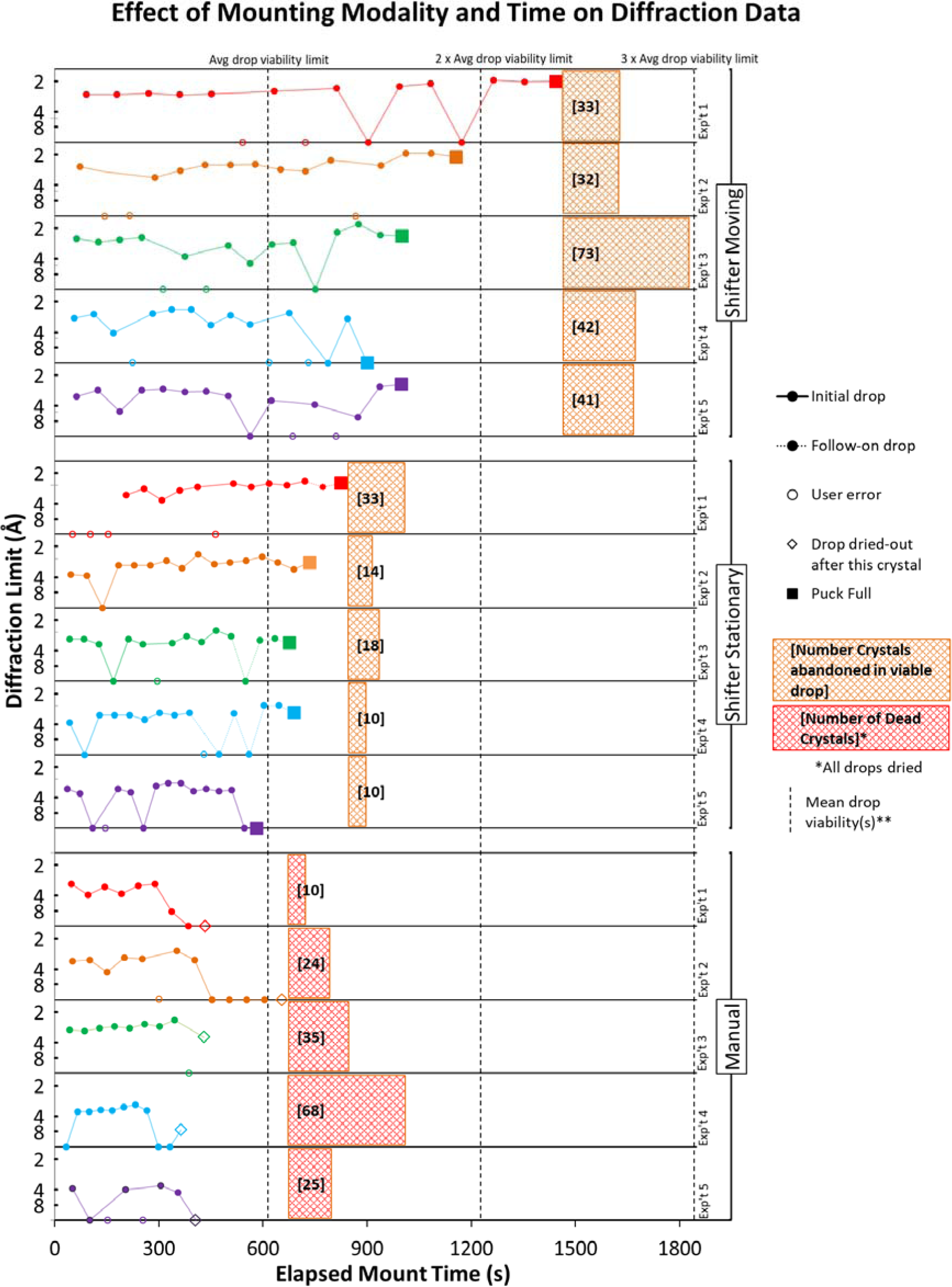
The Shifter enables mounting over a longer timeframe. The last crystal mounted from Shifter Stationary initial drops, before they became unusable, has a mean time stamp of 10.2 minutes. **Using this as a benchmark, dashed vertical lines indicate expected survival times for successive drops.

For Manually mounted wells, the drops dried with many crystals still present. In Shifter experiments more crystals are viable (Figure 8a-c) leading to more mounts, and more and higher resolution datasets (Figure 9). Though limited to 16 samples per experiment, many of the Shifter drops were still yielding viable crystals over timeframes long enough to have fully utilised all three drops. The hatched areas include first choice (35-75µm), and second choice (10-35µm) crystals left behind in viable drops (orange hatching) or dried drops (red hatching).

It should be emphasised that if a particular crystal system is difficult to mount from, then the Shifter will not in itself alleviate that specific problem (S6). Nevertheless, we have observed repeatedly that the improved ergonomics of provided by the Shifter appears to facilitate the same comparative improvement for all user experience levels and degrees of crystallisation system mounting difficulty.

### 3.4. High Positional Accuracy is Achieved Despite Low-Cost Components

We achieved a monotonic relationship between encoder value and stage position with acceptable positional accuracy from low-cost encoders. Errors in the accuracy of the position sensing system come primarily from non-linearity of the sensor (3% according to the manufacturer), apparently due to non-uniform thickness of the sensor. The evaluation of polynomial functions to find the best function for elucidation of real-world coordinates found that a 12th order function robustly generates well-distributed residual errors, with a positional accuracy of 0.1-0.15mm, over a range of representative calibration datasets (Figure 10).

**Figure 10.**
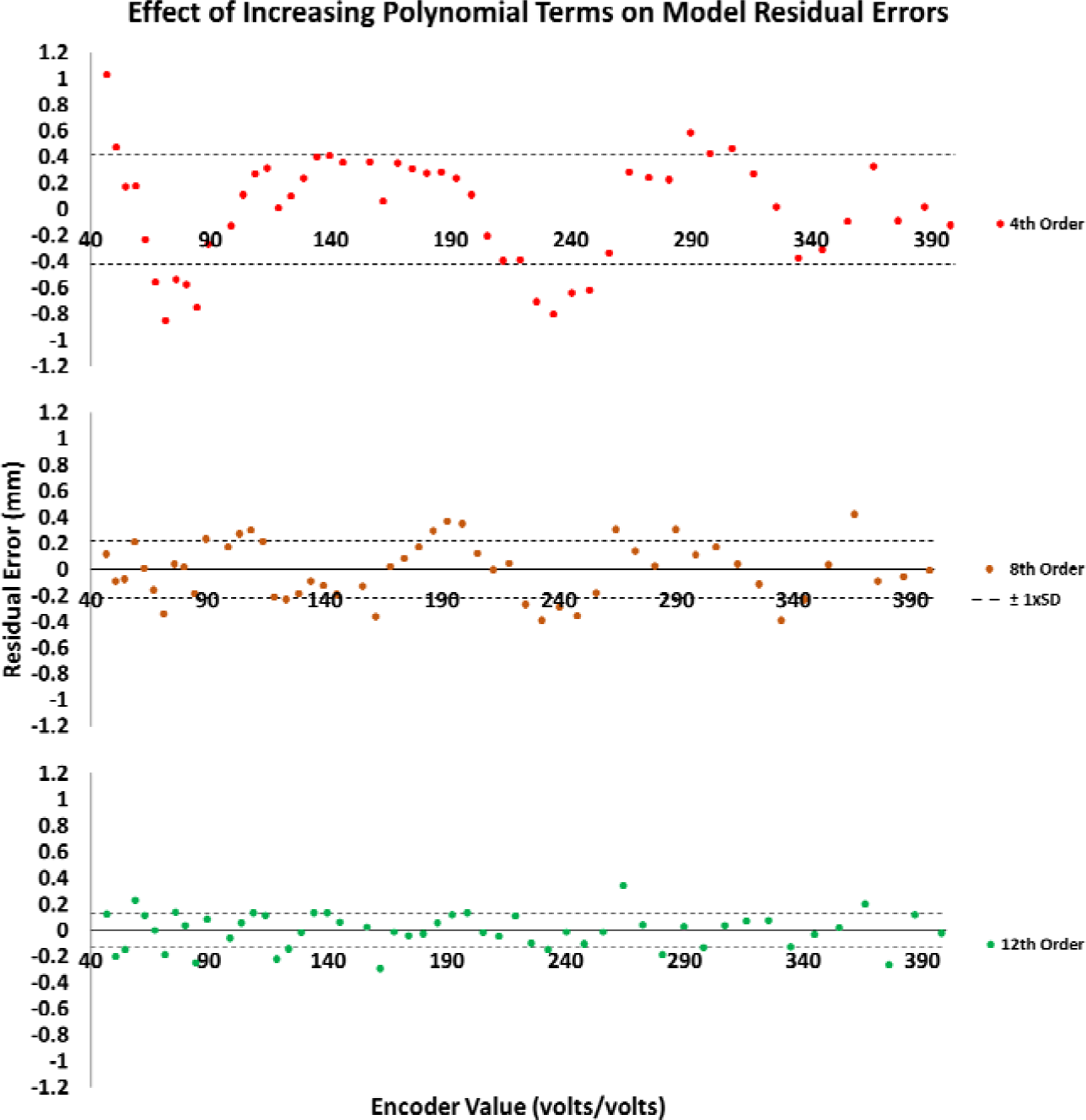
Accuracy can be achieved through calibration: This typical calibration dataset, fitted polynomial functions all have means for the error residuals not significantly different from zero. Residuals for lower (2-5) and middle (6-8) order polynomials usually fail the assumption of normality, higher (8-12) order polynomials show no trend over full length of the stage position encoder. As the numbers of orders included increases, the variation of the fitted model to the real-world coordinates becomes acceptable for the current application (4th, SD 0.4mm; 8th, SD 0.2mm, 12th, SD 0.1mm).

### 3.5. Future Work

The Shifter will be enabling for many more experiments than those reported here. To date, we have used the device only at room temperature, but the enclosure lends itself to cooling to low temperatures. This would accommodate crystal systems that must be maintained at 4°C, without moving the experiment to a cold room. Initial tests successfully maintained the temperature inside the enclosure below 0°C and above 90%(RH) for extended periods, without condensation internally or externally. The remaining challenge is in identifying a suitable air cooling and delivery system.

### 3.6. Lessons in hardware development for bench scientists

Through the open source and Maker movements, barriers to prototype engineering continue to fall, a trend first seen in industry in the 1970s (Augarten, 1974). This presents practitioners with great opportunities for innovation. With often free design software, end-users have access to tools to create bespoke apparatus at low-cost (Lian, Swainson, Cranswick, & Donaberger, 2009; J. M. Pearce, 2013). Popular microcontroller-powered sensing and control modules similarly allow designers to develop sophisticated systems, without in-depth electronics or software training. The economies of scale that come with a large customer base and active user community, mean they even make their way into low-volume commercial products (Cooke, 2017; Fryer, 2014).

However, hardware development remains a difficult and protracted process, if the goal set is simply to deliver a stable design that can be reproduced and operated independently by others. Designs must be thoroughly exposed to real-world use to uncover design limitations, and highlight where resource needs to be invested or can be omitted. This iterative and time-consuming process of ‘hardnening’ will reveal whether the intended audience sees adequate value in the proposed solution, and where the real user need lies. As in all areas of design, the technological road to quantity and quality of data in MX is paved with noble endeavours that have failed to achieve community penetration. Links to 3D print files or code repositories are not enough to enable the reproduction of a result in hardware development, as this requires expertise, and frequently an amount of development similar to designing from scratch. Since existing systems have already proven themselves to be robust, any attempt at addressing weakness in a process will struggle for adoption if it is not user-ready, or else wholly independent of a supporting infrastructure (Weissenberger, 2013), at least until performance is persuasive to the bulk of the user-base (Tellis, 2006).

## 4. Conclusions

The system described has proven itself to be an enabling solution, effectively addressing the sample preparation bottleneck. By integrating with and advancing current practices, users are not required to transition workflows to new, incompatible systems, in order to achieve productivity gains. The Shifter combines the aptitudes of humans and of machines, to provide a cost-effective, adaptable instrument where the inaccuracies inherent in low cost hardware choices are compensated for by software solutions and human skill. The need for complex engineering solutions to other design challenges has also been circumvented by careful consideration of the cost against its benefit.

The mounting bottleneck was a critical impediment to the XChem (DLS) workflow. By developing the Shifter in close collaboration with the XChem facility, this project has met current experimental needs, enabled new methodologies, and improved productivity for a range of internal and external users. The combination of machine semi-automation and auto-completion of tracking information has reduced time taken to fill a 16 sample puck from 60-80 minutes, down to 10 minutes. Mounting 100 crystals/h is now considered routine, with rates of ≥240 crystals/h achievable. The Shifter also alleviates the need for working in a cold room, when done purely to extend drop survival during mounting. The enclosure lid assists in organisation of hand tools and significantly steadies the hand by providing a resting surface for wrists. In the 4 years of XChem’s operation its users have mounted in excess of 100,000 crystals using the three Shifters installed.

Further to enabling high-throughput screening, Shifter sample protection and automated note-taking/data management prove useful even in single droplet work. The Shifter prevents noticeable evaporation during typical working times routinely exceeding 30minutes per plate. By extending droplet viability and reducing sample deterioration users are able to retrieve more crystals that embody information that would otherwise be lost. We predict that by increasing crystal yield and reducing crystal waste in this way there will be a reduction in protein purification and other upstream work. Whether from experimental or practical inefficiencies, the low productivity engendered by manual mounting translates ultimately into a higher ‘per outcome’ cost for the science generated (Adams, 2008; Pareek, Smoczynski, & Tretyn, 2011; Wetterstrand, 2016).

It was seen that the bottlenecks in mounting come as much from non-mounting tasks such as administration and seal cutting, as they do from crystal handling. The Shifter goes a considerable way to addressing these issues. Much of the success of this project has come from integrating the Shifter into user workflows, enabling a single user to carryout high throughput experiments where previously two people were needed. Though it was not a primary objective, GUI optimisation has been a key feature of our design solution (Leikanger, Balters, & Steinert, 2016).

The prototype described was developed at SGC and deployed to XChem (DLS) in 2015, where it remains in service. In 2016, a product based on the prototype was commercialised by one of the authors (NM).

## Acknowledgements

We thank the following SGC members, for testing and input: David Bowkett, Oakley Cox, Belinda Faust, Janine Gray, Jolanta Kopec, Tobias Krojer, Elizabeth MacLean, John Raynor, Laura Daiz Saez, Marion Schuller, Ritika Sethi, Rachael Skyner, Fiona Sorrell, Andrew Thompson and Eleanor Williams. From Diamond, the following helped with testing and integration: Jose Brandao-Neto, Alexandre Dias, Alice Douangamath and Renjie Zhang. David Jonathan Wright and Marine Peyret-Guzzon gave valuable input. At the Rutherford Appleton Laboratory (Harwell), David Wilsher, Sam Allum, and Andrew Lintern helped with additive manufacturing. From the University of Oxford, Department of Oncology, John Prentice and Gerald Shortland gave advice on machining, and Robert Newman and Iain Tullis provided useful discussions on electronics. The SGC is a registered charity (No. 1097737) that receives funds from AbbVie, Bayer, Boehringer Ingelheim, The Canada Foundation for Innovation, The Canadian Institutes for Health Research, Genome Canada, GlaxoSmithKline, Janssen, Lilly Canada, The Novartis Research Foundation, The Ontario Ministry of Economic Development and Innovation, Pfizer, Takeda and The Wellcome Trust (092809/Z/10/Z).

## Supporting information

### S1. Encoder Connectivity and Calibration

Encoder counts are related to real-world coordinated by comparing the encoder output to fixed scale markers.

**Figure 11.**
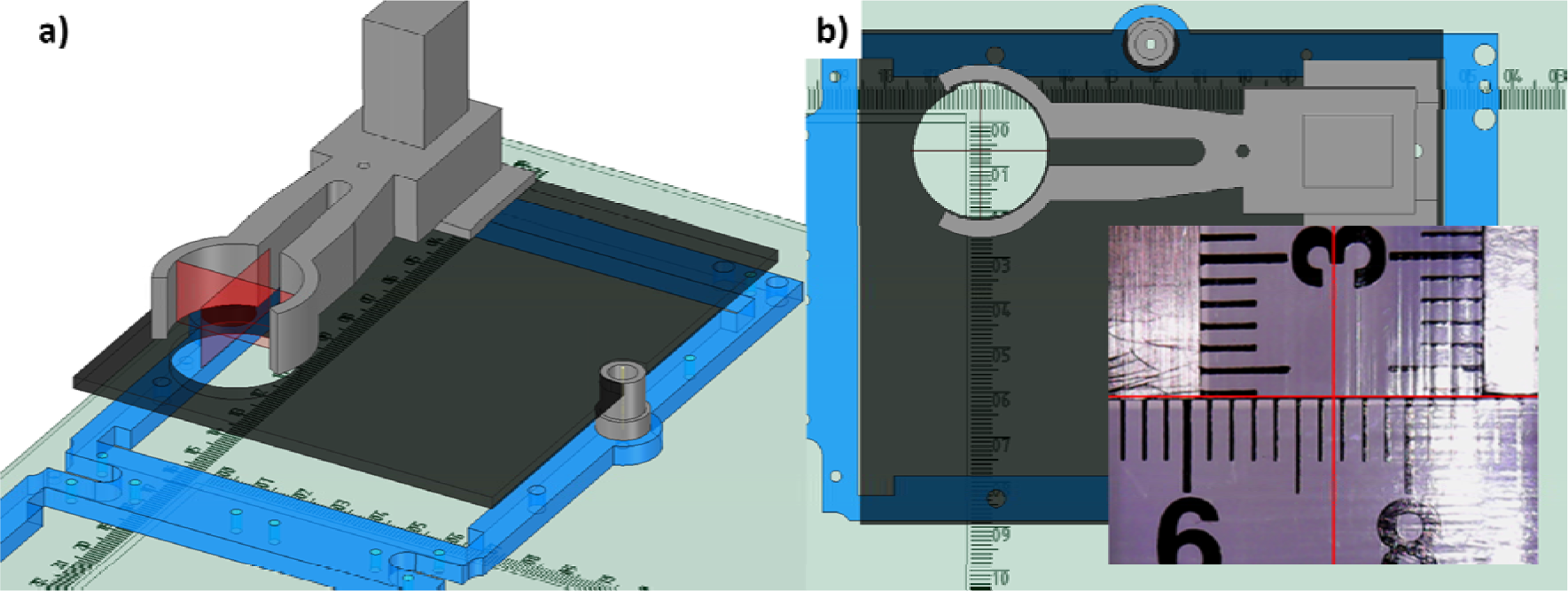
Calibrating the encoder: **a)** A USB microscope is clipped into a support (grey), which is fixed to a base (black). This assembly is mounted on the Y Carriage (blue). **b)** Cross-hairs (red) on the camera image overlay the X and Y scales on the enclosure base (green).

### S2. Survey of SGC Mounters on Time Spent Mounting

Six crystallographer responded to a self-response survey of time use and mounting practices as encountered in the community.

##### Survey Question

*Q: “How long does it usually take you to manually mount a puck of crystals (16 pins)?”*

*A: Preparation/Pre-mounting (preparing mounting lists, etc.)_____________________________*

*A: Mounting (all related processes; cutting seals, finding wells, making notes etc.) ____________*

*A: Post-mounting (transcribing notes, registering in database, etc.)____________*

**Table 2.** Time used per phase of mounting process (minutes)

| User | Preparation | Mounting | Post-mounting | Total | Time ‘off task’ (%) |
| --- | --- | --- | --- | --- | --- |
| 1 | 10 | 60 | 10 | 80 | 25 |
| 2 | 45 | 120 | 60 | 225 | 47 |
| 3 | 12 | 50 | 15 | 77 | 35 |
| 4 | 20 | 40 | 15 | 75 | 47 |
| 5 | 30 | 60 | 30 | 120 | 50 |
| 6 | 20 | 150 | 20 | 190 | 21 |

Manual mounting is slow and distracted. Self-reported data show long mount times (M: 8min, SD: 4min) of 7.5 crystals/h, with 38% of time devoted to non-mounting tasks.

### S3. XChem User Data: September 2015 and January 2016

Crystal mounting outcomes captured via GUI button press during user operation. ‘Mounted_x’ indicates a successful mount followed by a descriptor of the drop appearance; ‘Mounted_Clear’ is the typical mode for optimised systems.

**Table 3.** Time used per phase of mounting process (minutes)

| Outcome Mode | Number of Mounts | (%) |
| --- | --- | --- |
| Fail_Evaporated | 66 | 1 |
| Fail_Melted | 933 | 11 |
| Fail_Unpickable | 150 | 2 |
| Mounted_BadDispense | 135 | 2 |
| Mounted_Clear | 6112 | 74 |
| Mounted_Crystalline | 320 | 4 |
| Mounted_Precipitate | 555 | 7 |
| All Failures | 1149 | 14 |
| All Successes | 7122 | 86 |
| Total | 8271 |  |

**Figure 12.**
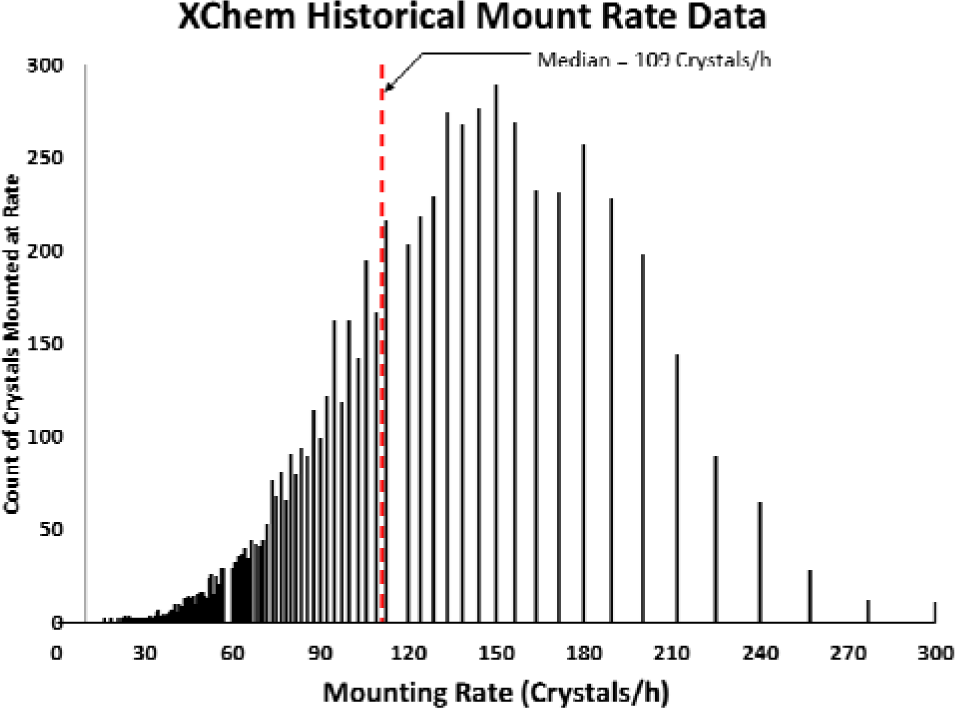
Mount rates of users from the same dataset as Table 3. The mount duration for each mounted crystal was recorded in whole seconds, and converted to an hourly rate with the following formula:

### S4. Droplet humidification techniques

Evaluation of bubble-column humidifier configurations, against the practice of using a domestic ultrasonic humidifier.

**Table 4.**

| Humidification Type | Reservoir Solution | Reservoir Temp. (°C) | Configuration (Column depth or power setting) | Protected Atmosphere Humidity (%RH) | Humidified Air Temp. (°C) |
| --- | --- | --- | --- | --- | --- |
| Bubble | Water | 4 | 70ml | 42 | 21 |
| Bubble | Water | 83 | 70ml | 96 | 30 |
| Bubble | 5M NaCl | 20 | 70ml | 65 | 20 |
| Bubble | 1.5M NaCl | 20 | 70ml | 70 | 20 |
| Bubble | Water | 20 | 70ml | 74 | 20 |
| Bubble | Water | 20 | 140ml | 98 | 20 |
| Ultrasonic* | Water | 20 | Min. power | 97 | 20 |
| Ultrasonic* | Water | 20 | 60% power | 98 | 20 |

\*ultrasonic humidifier at the work area.

**Figure 13.**
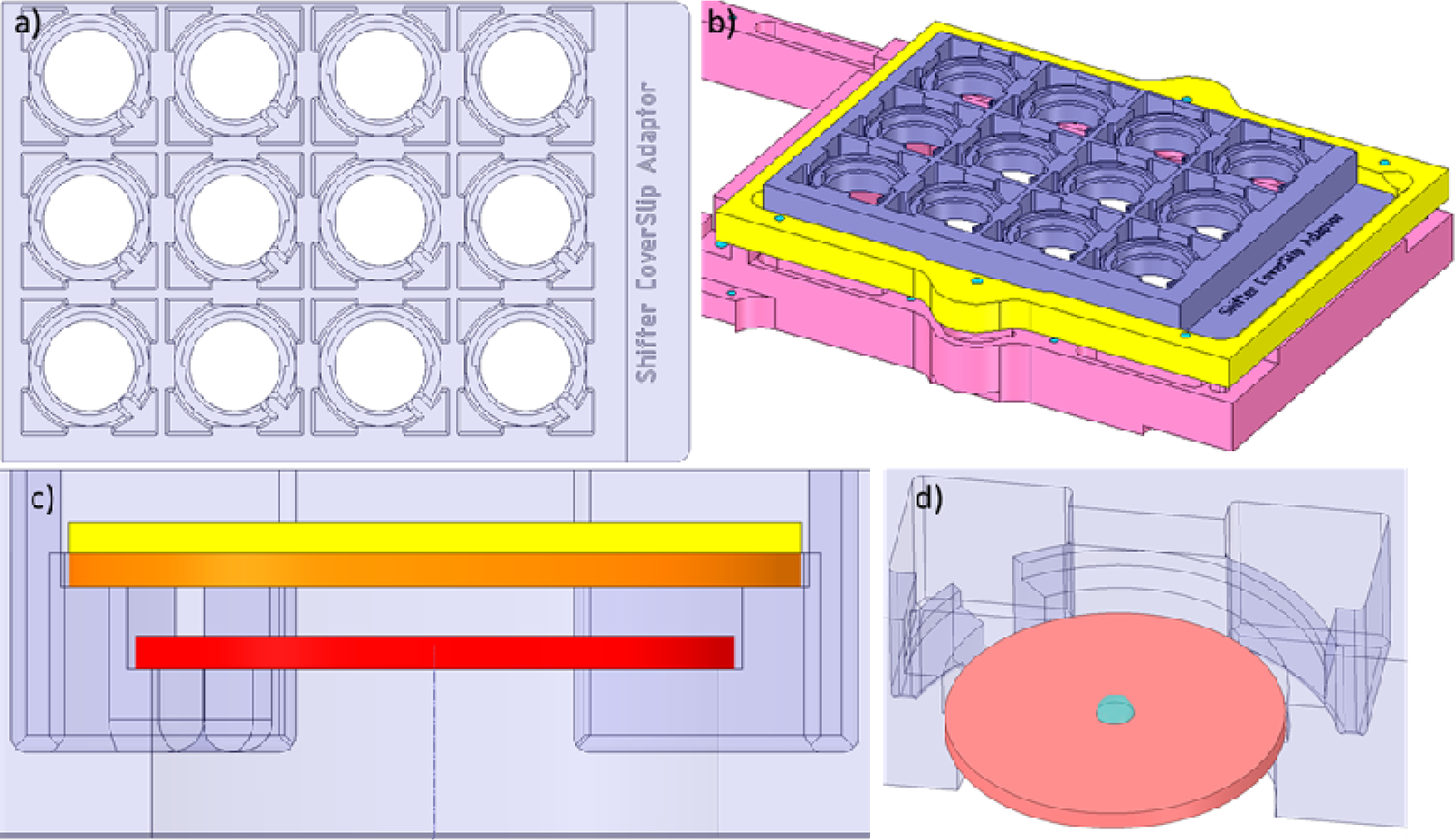
Cover slip adaptor for inverted hanging-droplet experiments **a)** plan view, **b)** trimetric view, **c)** Cross-section and **d)** cutaway, showing position of 18mm (red) and 22mm (orange) round, and 22mm square (yellow) coverslips.

**Figure 14.**
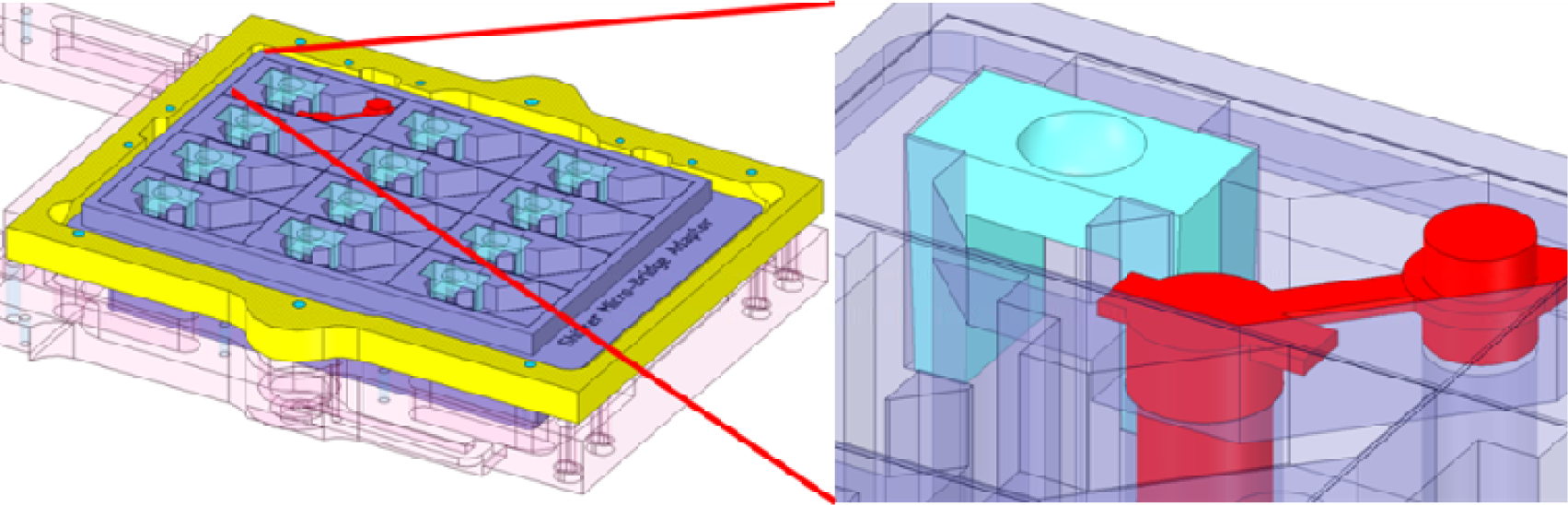
Micro-Bridge (light blue) adaptor with removable 0.2ml PCR tube reservoir (red)

### S6. Mounting from difficult drops remains somewhat difficult

An experienced mounter (RT) harvested from a selection of crystal systems judged to be representative of differing mounting difficulty: BRD1A (easy mounting), DACASA (moderately easy), and JMJD1BA (difficult mounting). Median mounting times were 25, 20 and 29 seconds per crystal respectively (124-180 crystals/h)(Figure 15). Whilst there is no trend between ‘mounting ease’ and absolute speed of mounting in this experiment, mounting does seem to become less skewed as the difficulty of retrieving crystals increases. Mount times of 13-15 seconds per crystal seem to represent the practical limit of the mounting process, not because of the speed of the Shifter (move times 1-3 seconds are typical), but because of mounting, freezing and storing the crystal, and preparing for the next mount. Whilst the Shifter cannot ‘cure’ the effect of a difficult system on mounting speed, metrics captured by the system now allow for routine analysis of such trends.

**Figure 15.**
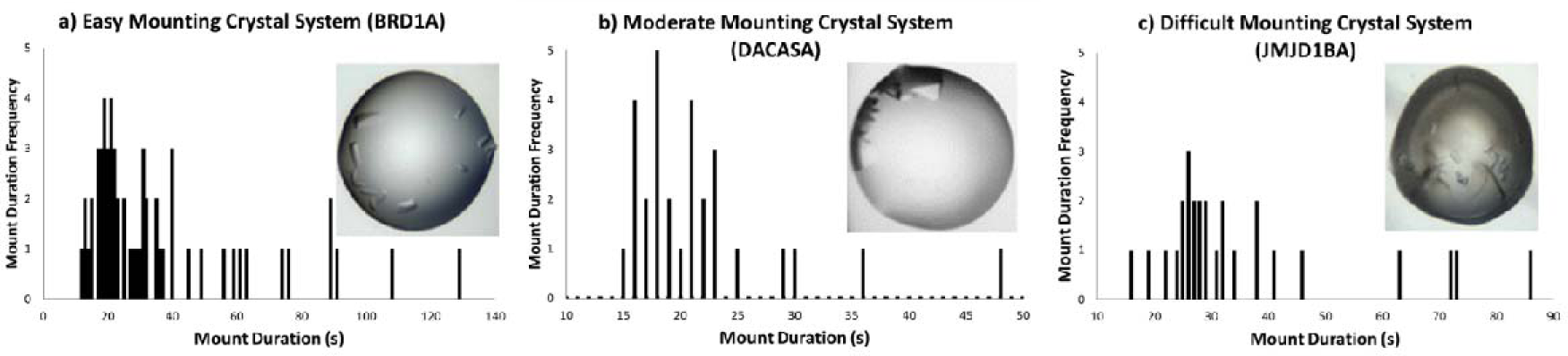
Crystal mount time distributions: a) BRD1A n=59, median 25s; b) DACASA n=29, median 20s; c) JMJD1BA n=27, median 29s

